# Lipoprotein particles interact with membranes and transfer their cargo without receptors

**DOI:** 10.1101/2020.08.27.270496

**Authors:** Birgit Plochberger, Taras Sych, Florian Weber, Jiri Novacek, Markus Axmann, Herbert Stangl, Erdinc Sezgin

**Affiliations:** TU Wien, Institute of Applied Physics, Vienna, 1040, Austria; Johannes Kepler University Linz, Institute of Biophysics, Linz, 4020, Austria; Upper Austria University of Applied Sciences, Campus Linz, Linz, 4020, Austria; Science for Life Laboratory, Department of Women’s and Children’s Health, Karolinska Institutet, 17165, Solna, Sweden; CEITEC, Masaryk University, University Campus Bohunice, Brno, 62500,Czech Republic; Medical University of Vienna, Center for Pathobiochemistry and Genetics, Institute of Medical Chemistry, Vienna, 1090, Austria; MRC Human Immunology Unit, MRC Weatherall Institute of Molecular Medicine, University of Oxford, Oxford, OX3 9DS, UK

**Author notes:** Equal contribution (Birgit Plochberger, Taras Sych). **Correspondence: Erdinc Sezgin** 00441865222484 **Birgit Plochberger**: 00435080452131.

**Keywords:** HDL, LDL, VLDL, cholesterol, receptor-independent cargo transfer, LUV, GUV, GPMV, SLB, fluorescence correlation spectroscopy, spectral imaging, generalized polarization

## Abstract

Lipid transfer from lipoprotein particles to cells is essential for lipid homeostasis. High density lipoprotein (HDL) particles are mainly captured by cell-membrane-associated scavenger receptor class B type 1 (SR-B1) from the blood stream while low and very low density lipoprotein (LDL, VLDL) particles are mostly taken up by receptor-mediated endocytosis. However, the role of the target lipid membrane itself in the transfer process has been largely neglected so far. Here, we study how lipoprotein particles (HDL, LDL and VLDL) interact with synthetic lipid bilayers and cell-derived membranes and transfer their cargo subsequently. Employing cryo-electron microscopy, spectral imaging and fluorescence (cross) correlation spectroscopy allowed us to observe integration of all major types of lipoprotein particles into the membrane and delivery of their cargo in a receptor-independent manner. Importantly, biophysical properties of the target cell membranes change upon cargo delivery. The concept of receptor-independent interaction of lipoprotein particles with membranes helps to better understand lipoprotein particle biology and can be exploited for novel treatments of dyslipidemia diseases.

## Introduction

Cholesterol is a major structural element in cell membranes (1). Thus, a steady supply of cholesterol is of utmost importance for cell membrane integrity. Its levels are tightly controlled by homeostatic mechanisms balancing pathways of cholesterol uptake, biosynthesis and release (2). Specialized cargo vehicles, called lipoprotein particles, are necessary to solubilize their share of non-polar cargo. Several pathways are operative for cholesterol uptake – the majority via receptor-mediated endocytosis, in which low density lipoprotein (LDL) receptor binds apoB- and apoE-containing lipoprotein particles (3,4) which are subsequently endocytosed. In addition, selective lipid uptake via Scavenger Receptor Class B family (SR-B) receptors, in which core lipids of the lipoprotein particles are transferred to cells and tissues, has been proposed (5). Furthermore, endocytosis and subsequent transcytosis of lipoproteins particles is operative at least in endothelial cells (6, 7). Besides these well-described pathways, direct transfer of cholesterol from lipoprotein particles to cell membrane may also exist (8–12). Here, using advanced imaging techniques, we show that cholesterol is transferred from all lipoprotein particles to lipid-only large and giant unilamellar vesicles (LUVs, GUVs), supported lipid bilayers (SLBs) as well as cell-derived giant plasma membrane vesicles (GPMVs). Upon delivery, rigidity of the target membrane increases as expected due to stiffening effect of cholesterol.

## Materials and Methods

### Reagents

Alexa Flour 647 NHS ester was obtained from Invitrogen. Sephadex G-25 fine resin, sodium cyanoborohydride (NaCNBH3), TriEthylAmine (TEA), 3-AminoPropyl-TriEthoxySilan (APTES), EthanolAmine (ETA), sodium deoxycholate, sucrose, glucose and HEPES were from Sigma. 1-palmitoyl-2-oleoyl-sn-glycero-3-phosphocholine (POPC) and cholesterol linked to BodipyFL (TopFluor-Cholesterol, Bd-Chol) was obtained from Avanti Polar Lipids. C-Laurdan was purchased from 2pprobes (2pprobes.com). NR12S was provided by Dr. Andrey Klymchenko, University of Strasbourg. Abberior Star Red DOPE was purchased from Abberior.

### Lipoprotein particle isolation and labeling

Blood donations, obtained from normolipidemic healthy volunteers, were approved by the Ethics Committee, Medical University of Vienna (EK-Nr. 511/2007, EK-Nr. 1414/2016). Lipoprotein particles were isolated as previously described (8) via sequential flotation ultracentrifugation. Its proteins were covalently linked to Alexa Fluor 647 at pH=8.3 according to the manufacturer’s instruction. Bd-Chol was incorporated into the lipid leaflet of lipoprotein particles via incubation at 37 °C for 2 h. Free dye and excessive cholesterol was removed via extensive dialysis.

### Preparation of giant unilamellar vesicles (GUVs)

GUVs with varying sizes from 10 μm to 100 μm were prepared by electroformation (9, 10). POPC was dissolved in chloroform (1 mg/ml) and deposited on Pt electrodes. The solvent was evaporated by a constant N_2_ flow for 20 min. 370 μl of 300 mM sucrose was added in a self-made chamber. On the cap of this chamber, we placed two holes with a distance of 5 mm for the electrodes. After, the electrodes with dried lipids were incubated in the sucrose solution, a voltage of 2 V at 10 Hz for 1 h and for another 30 min at 2 Hz was applied at room temperature (≈23°C). Fluorescently labeled lipoprotein (HDL, LDL or VLDL) solution was added to the GUV solution (final concentration of lipoproteins 0.3 mg/ml). Images were acquired 20 min after addition by confocal microscopy. For imaging, we added 100 μl of GUV suspension in sucrose to 100 μl of PBS.

### Preparation of large unilamellar vesicles (LUVs)

LUVs were prepared by extrusion (Avanti Mini Extruder, Avanti Polar Lipids, USA). DOPC was dissolved in chloroform/methanol (2:1, 10 mg/ml) and 10 μl were dried by evaporation. Subsequently, lipids were hydrated using PBS buffer and kept above the phase transition temperature of the lipid during hydration and extrusion. Once the sample is fully hydrated, the mixture was placed into one end of the Mini-Extruder. The plunger of the filled syringe was pushed gently until the lipid solution is completely transferred to the alternate syringe and afterwards the plunger of the alternate syringe was pushed to transfer the solution back to the original syringe. This process was repeated until the lipid suspension was clear. The lipid solution was stored at 4°C.

### Preparation of giant plasma membrane vesicles (GPMVs)

CHO cells were grown in DMEM/F12 medium supplemented with 10% FBS and 1% L-glutamine. GPMVs were prepared as previously described (10). Briefly, cells seeded out on a 35 mm petri dish (≈70 % confluent) were washed with GPMV buffer (150 mM NaCl, 10 mM Hepes, 2 mM CaCl_2_, pH 7.4) twice. 1 ml of GPMV buffer was added to the cells. 25 mM Paraformaldehyde and 2 mM Dithiothretiol (final concentrations) were added in the GPMV buffer. Cells were incubated for 2 h at 37 °C. Then, GPMVs were collected by pipetting out the supernatant.

### Preparation of supported lipid bilayers (SLBs) on mica substrate (for AFM Force Spectroscopy)

SLBs were formed on freshly cleaved (~1 μm thick) mica, glued onto a glass coverslip using an optically transparent UV-glue (optical adhesive 88, Norland Products Inc.). Glass slides (d = 22 mm, Menzel) were incubated in a freshly prepared mixture of sulfuric acid: hydrogen peroxide (3:1) for 20 min, rinsed with deionised water and ethanol and dried under N_2_ flow. 30 μl of DOPC solution (10 mg/ml in chloroform: methanol, 3:1) was evaporated under N_2_ flow (20 min) and resuspended in 300 μl PBS. Vesicles were prepared by sonication for 20 min and applied to the mica. After 20 min, the bilayer was formed and slides were washed with PBS.

### Preparation of supported lipid bilayers (SLBs) on glass substrate (for fluorescence microscopy)

POPC was dissolved in chloroform/methanol (2:1, 1 mg/ml final concentration) with or without 0.01 mol % of Abberior Star Red DOPE. Glass coverslips (25 mm in diameter, #1.5) were incubated in a freshly prepared mixture of sulfuric acid:hydrogen peroxide (3:1) for 30 min, rinsed with deionised water and dried under N_2_ flow. Glass coverslip was mounted on the spin-coater (SPI supplies), spinning started at 3000 rpm, 25 μl of lipid solution was applied and spinning was continued for 30 seconds. The glass coverslip was immediately mounted on the metal Attofluor Cell Chamber and the lipid film was hydrated with the SLB buffer (150 mM NaCl, 10 mM HEPES, pH=7.4).

### Confocal Microscopy

GUVs and GPMVs were imaged with a laser scanning confocal microscope (LSM 780, Zeiss). The microscope was equipped with a 40x/1.20 water immersion objective. 488 nm and 633 nm lasers were used to excite BodipyFL and Alexa Fluor 647, respectively. Spectral imaging of C-Laurdan and NR12S was performed on a Zeiss LSM 780 confocal microscope equipped with a 32-channel GaAsP detector array. Laser light at 405 nm was used for fluorescence excitation of Laurdan. The lambda detection range was set between 415 nm and 691 nm for Laurdan. Laser light at 488 nm was used for fluorescence excitation of NR12S. The lambda detection range was set between 498 nm and 691 nm for NR12S. Images were saved in .lsm file format and then analyzed by using a freely available plug-in compatible with Fiji/ImageJ, as described (11).

### Force Spectroscopy and Bilayer Indentation Experiments

Force measurements were performed on a PicoPlus AFM (Agilent Technologies) operated under PicoView 1.6.8 (Agilent Technologies) in solution (PBS Puffer). Force distance cycles were acquired using silicon cantilevers with a spring constant of 0.01 N/m or 0.02 N/m (Veeco) at pulling velocities of 0.1 μm/s – 5 μm/s and contact times (hold times) between 0.1 – 5 seconds. Empirical force distributions of the rupture forces of the last unbinding event (PDF) were calculated as described (12). PDFs were fitted with the equation 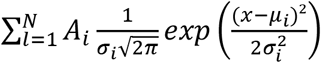 including the boundary condition 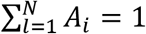 taking the probability density property of the PDFs into account, where *A*_*i*_ is a prefactor, *μ*_*i*_ is the position of the peak, and *σ*_*i*_ is the width of the peak (see Fig. S1). For force spectroscopy, a sweep range of 3 μm and sweep rate of 0.2 - 2 Hz were used. Silicon-nitride AFM cantilevers with silicon tips (MSNL-10, Bruker AFM Probes) were amine-functionalized as described previously (13). Briefly, silicon cantilevers were amine-functionalized via gas-phase silanization with AminoPropyl-TriEthoxySilane (APTES) (14) and a heterobifunctional (aldehyde-NHS) linker was chemically connected. Subsequently, the tips were washed with chloroform and dried with N_2_ gas. Tips were incubated with 100 μL lipoproteins (0.06 mg/mL in PBS), to which 2 μL NaCNBH_3_ (1 M, freshly prepared in 10 mM NaOH) was added for irreversible binding. Afterwards, 5 μL of ethanolamine hydrochloride (1 M, adjusted to pH 9.6) was added for blocking non-reacted linker groups and incubation was continued for 10 min. This chemical modification was used to covalently link lipoprotein particles on cantilevers. The effective spring constant was determined via thermal noise analysis (15) before and after chemical modification.

### AFM imaging and particle analysis

AFM measurements were performed with an Atomic Force Microscope (JPK BioAFM - NaonWizard 4, JPK, Berlin). AFM probes made of silicon nitride with nominal spring constant of 0.3 N/m and nominal tip radius between 20 nm – 60 nm (MLCT-BIO-F, Bruker Nano Inc., Camarillo, CA) were used for the measurements. The exact sensitivity and spring constant of each cantilever was determined on a cleaned coverslip in 300 μl PBS from a force-displacement experiment and a thermal noise spectrum measurement. All samples (HDL, LDL, VLDL) were diluted to 1:1000. A volume of 300 μl of the diluted HDL and LDL solution was incubated on the cleaned glass coverslip for at least 5 min and subsequently imaged. Because of the low density of VLDL particles a volume of 30 μl of the diluted VLDL sample was incubated on the glass cover slip upside down for 5 min. Afterwards AFM images were obtained by using an advanced imaging software (Quantitative Imaging mode QI™-mode) of Bruker. A maximal set point force of 500 pN was used.

Particle analysis (Full Width at Half Maximum (FWHM) and height of probe molecules) was performed with JPK Data Processing software (V.6.1.163, JPK, Berlin). Convolutions of tip artifacts were corrected as described in supplementary information (Fig. S2). For each individual particle, the aspect ratio ([AR] = %) was calculated. An AR of 100% represents a perfect spherical shape, lower values represent prone discs (Fig. S2).

### Cryo-electron microscopy

LUVs (100 μl) were incubated with respective lipoprotein solutions (5 μl) for 2 min at room temperature. Immediately after incubation, samples were stored on ice and applied to the cryo-grids (2 nm precoated Quantifoil R3/3 holey carbon supported grids) at a concentration of 10 μM and vitrified using a Vitrobot Mark IV (FEI). Data collections were performed on TEM microscope FEI Tecnai F20 equipped with a 4k CCD camera and two side-entry cryo-holders. Dataset was collected using Tecnai F20 (FEI, Eindhoven) operated at 200 kV and equipped with a 4k charge-coupled device detector FEI Eagle. Micrographs were collected with a pixel size of 1.79 Å and total dose of 20 e^−^/Å^2^. Frames were aligned using MotionCor2 (16), and CTF parameters were estimated using Gctf program (17).

## Results and Discussions

First, we imaged lipoprotein particles with atomic force microscopy (AFM) which confirmed intact particles (Fig. 1A). For this purpose, we incubated lipoprotein particles on clean glass for immobilization and performed AFM measurements. From these measurements, we calculated lateral and axial size for each particle (Fig. 1B-D; Fig. S2, 3). The sizes (lateral x axial) were 9.4 ± 2.1 nm × 8.3 ± 3.0 nm (aspect ratio (AR) = 92%) for HDL particles, 28.8 ± 8.7 nm × 22.5 ± 4.3 nm (AR = 84%) for LDL particles and 64.6 ± 5.1 nm x 48.8 ± 6.0 nm (AR = 76%) for VLDL particles in line with literature values obtained with other techniques (18). Fig. 1D shows the spherical reconstruction of the particles according to the size calculations from AFM images. After confirmation of particle structure and integrity, we set out to study the interaction of lipoprotein particles with membranes.

**Figure 1.**
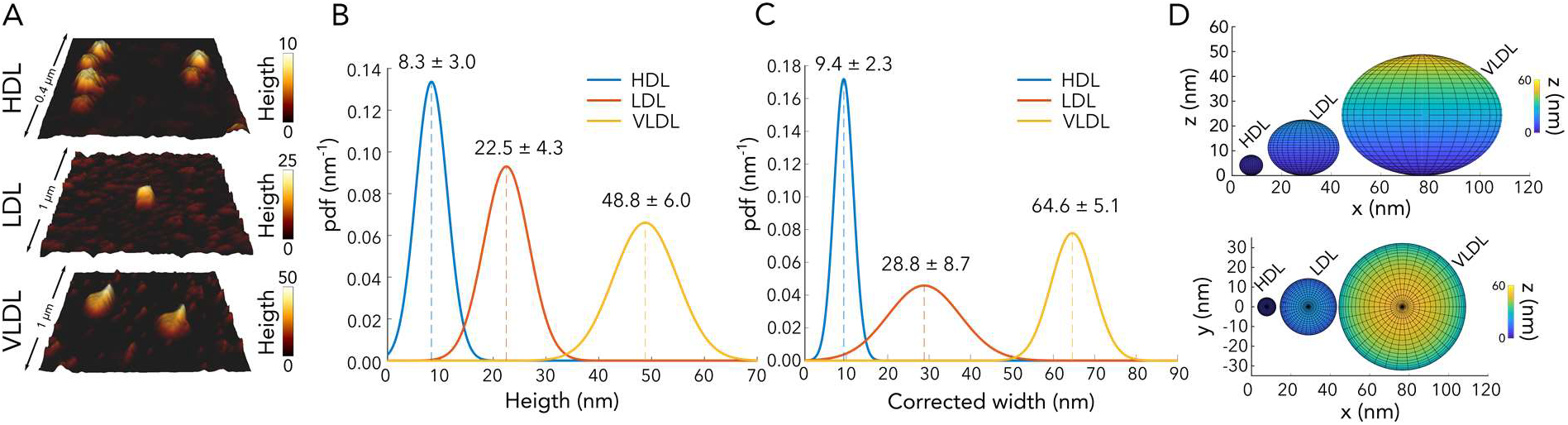
Topographical characterization of the lipoprotein particles with AFM. A) AFM images of lipoprotein particles. B) Height, C) width distribution (Mean ± Std) of 10 analyzed lipoprotein particles. D) Spherical reconstruction of lipoprotein particles according to the AFM data.

Recently, by using cryo-EM, we studied LDL interaction with membranes (19). This prompted us to study whether similar interaction pattern exists with all lipoprotein particles and with lipid-only membranes. We applied lipoprotein particles to large unilamellar vesicles (LUVs) and observed clear interactions between the lipoproteins and the LUVs (Fig. 2A). Interaction of different lipoprotein particles with LUV membranes (blue circles) was confirmed through recording data under different electron-beam incident angles, thus excluding an accidental overlay of signals originating from different layers of the vitrified ice.

**Figure 2:**
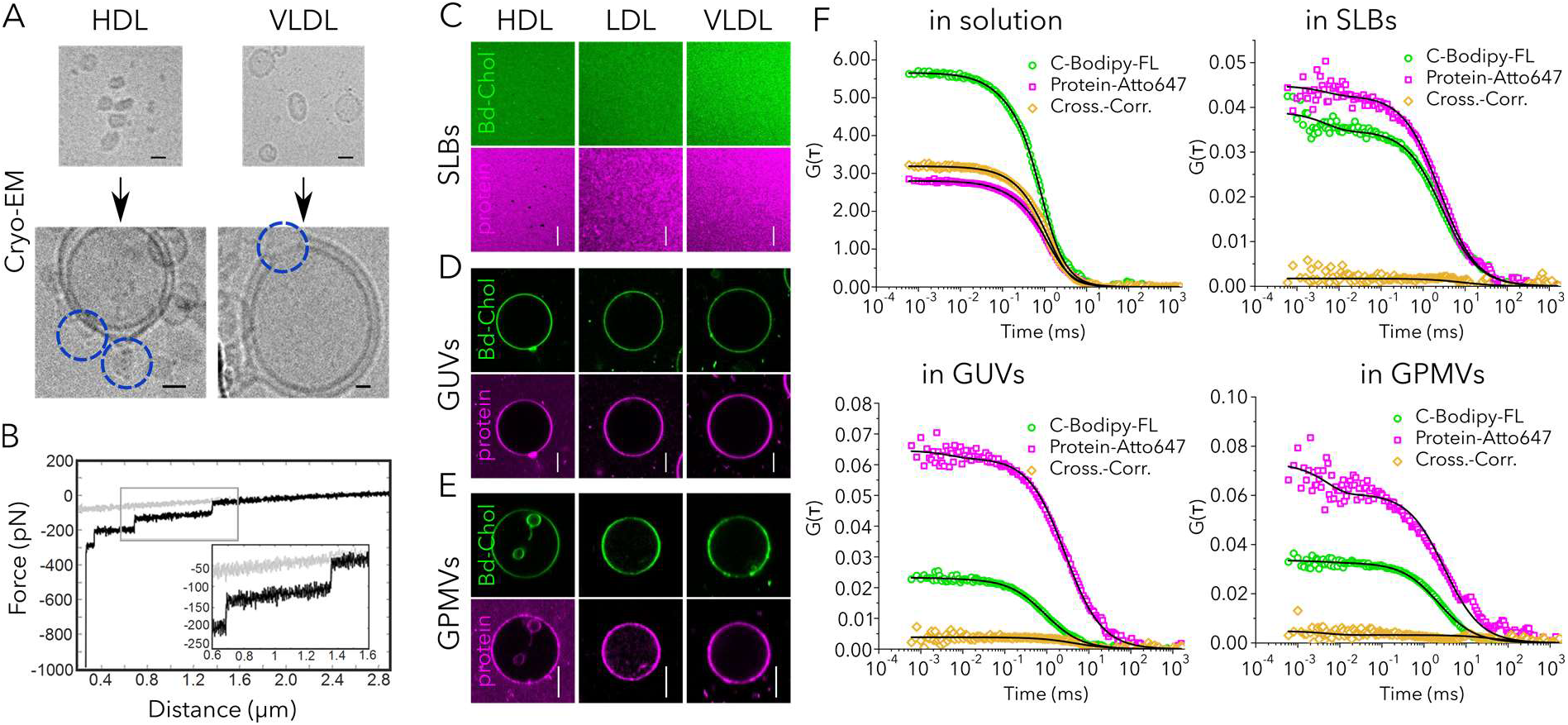
Lipoprotein particles interact with biomimetic membranes and transfer their cargo. A) Cryo-EM images of single HDL and VLDL particles (upper images) and lipoprotein decorated LUVs (lower images). Images were acquired under low-dose conditions (20e/Å^2^). Scale bar = 10 nm. B) Membrane tethers are formed during the retraction of HDL-modified AFM tips from supported lipid bilayers. A representative retraction curve (black) is shown for a functionalized HDL tip on a DOPC membrane. A cantilever with a spring constant of 0.01 N/m was used. The applied pulling velocity was 1 μm/s. For experiments, a maximum contact force of 500 pN was set in order to prevent penetration of the membrane. During retraction, membrane tethers are formed between the HDL particle on the tip and the bilayer with typical rupture forces of ~50 pN. Confocal images of C) SLBs D) GUVs and E) GPMVs incubated with 0.05 mg/ml fluorescently labelled HDL (upper images), LDL (middle images) and VLDL (lower images). Bd-Chol is depicted in green and Alexa 647 labeled protein is shown in magenta. Scale bars = 10 μm. F) Fluorescence cross-correlation spectroscopy of Bd-Chol and proteins measured in solution (intact VLDL) as well as in target membranes (GUVs, GPMVs and SLBs; see Fig S5 for HDL and LDL). In solution, high cross-correlation of Bd-Chol and protein signals is detected which suggests co-diffusion. In target membrane, cross-correlation curve amplitude is close to zero which suggests that Bd-Chol and protein molecules diffuse in the target membrane independently.

To further investigate the interaction of lipoprotein particles with membranes, we recorded AFM force-distance cycles. This technique is conventionally used to determine the interaction between molecular compounds (e.g. ligand-receptor). For this purpose, one binding partner is chemically bound to the AFM tip and the other is immobilized on the surface to be investigated or is located directly in the cell membrane. By bringing the tip to the surface, the binding of the molecules to be investigated is enabled, by pulling back the binding is broken, and the strength and kinetics of the interaction can be determined. In our case, lipid-only the membrane was used as binding partner of lipoproteins which was attached to the AFM tip. Specifically, HDL-functionalized silicon tips were brought in contact with supported lipid bilayers. The force setpoint was set below an actual penetration of the membrane and therefore the system experiences only a small counterforce. Interestingly, by performing force distance cycles with an HDL tip on a fluid supported lipid bilayer, interaction forces were detected characteristic for tube formation (see Fig. 2B, Fig. S4).

Next, to directly visualize whether interaction of lipoprotein particles lead to cargo transfer from VLDL, LDL and HDL particles to biomembrane systems, we applied fluorescence imaging. We used SLBs as supported and GUVs as free-standing lipid-only membrane systems. We labeled lipoprotein particles with Cholesterol-BodipyFL (Bd-Chol) and its proteins (Apolipoportein A for HDL and Apolipoprotein B for LDL) with Alexa Fluor 647. After incubation, we detected the fluorescence signal of both cholesterol and proteins in SLBs and GUVs for all types of lipoproteins (Fig. 2C, D). This suggests that upon interaction with the target membrane, lipoprotein particles transfer their cargo. To verify this observation in a more complex membrane system, we prepared GPMVs from Chinese Hamster Ovary (CHO) or HeLa cells, which comprise a biological membrane system consisting not only of lipids but also of proteins. Similarly, we observed cholesterol and protein transfer from lipoprotein particles to the GPMV membrane (Fig. 2E). To unequivocally verify the cargo transfer, we applied fluorescence cross correlation spectroscopy (FCCS). FCCS is a fluctuation-based method that measures the co-diffusion of two different fluorescently labelled molecules (20). It yields an autocorrelation curve for both differently labelled molecules providing information on their diffusion, concentration and molecular brightness. Additionally, it yields a non-zero amplitude cross-correlation curve if molecules co-diffuse. Co-diffusion usually means direct interaction or association to the same nanoscale entity (such as a domain, vesicles etc.) that moves through the focal spot. The amplitude of the cross-correlation curve is proportional to the co-diffusion rate; perfect co-diffusion gives near 100% while no co-diffusion yields 0% cross-correlation. In principle, intact lipoprotein particles where cholesterol and proteins are both labelled should show perfect co-diffusion with high amplitude as both molecules move in and out of the focal volume together as one particle unit. We indeed observe very high cross correlation (nearly 100%) for all types of lipoprotein particles in solution (Fig. 2F, Fig. S5). If the content of the lipoprotein particles is released into their target membrane upon interactions, fluorescently-labelled proteins and cholesterols should move separately in the membrane (unlike in solution). Thus, cross correlation should disappear. To test this, we measured the cross-correlation in SLBs, GUVs and GPMVs incubated with labeled lipoprotein particles and indeed we observed no cross correlation in these samples (near 0 amplitude, Fig. 2F, Fig. S5). Moreover, in solution, diffusion coefficient of cholesterol and protein is identical since they move together in the same particle. However, once lipoproteins fuse with the target membranes, diffusion of protein is always slower than cholesterol diffusion due to its size (Fig. S6). This data confirm that lipoprotein particles interact with the synthetic membranes and subsequently release their cargo to the target membranes.

Cholesterol content in the plasma membrane is crucial for membrane biophysical properties such as rigidity, stiffness, elasticity etc. Therefore, it is necessary to reveal how interaction and cargo transfer of lipoprotein particles alter biophysical properties of target membranes. We measured the rigidity of target membranes by using two environment-sensitive probes NR12S (21) and C-Laurdan (22). The fluorescence emission of these probes is sensitive to the rigidity of the lipid environment; they demonstrate red shift in their emission maximum in more fluid membranes. This spectral shift can be used to report on the molecular ordering of the membranes, utilizing an empirical lipid packing parameter, generalized polarization (GP) (23). GP in an indirect but robust way to infer lipid packing with values varying between +1 (for very ordered) and −1 (very disordered) (24). Confocal spectral imaging can conveniently be used to measure GP values of membranes (11). To measure the GP values of membranes, we incorporated NR12S in GUVs, C-Laurdan in GPMVs and imaged them with spectral imaging before and after incubation with lipoprotein particles. Higher cholesterol content yields more rigid membranes, thus higher GP values. While control GUVs (not treated with any lipoprotein particles) yielded GP values of −0.24±0.07, GUVs incubated with lipoprotein particles showed GP values of −0.19±0.05 (HDL), −0.08±0.08 (LDL) and −0.1± 0.07(VLDL) (Fig. 3A, B). Similarly, GPMVs that are not treated with lipoprotein particles showed −0.07±0.02 while lipoprotein-treated ones showed 0.05±0.02 (HDL), 0.04±0.01 (LDL) and 0.0±0.01 (VLDL) (Fig. 3C, D). This data shows that upon cargo transfer, cholesterol is incorporated in the membrane and rigidifies it.

**Figure 3.**
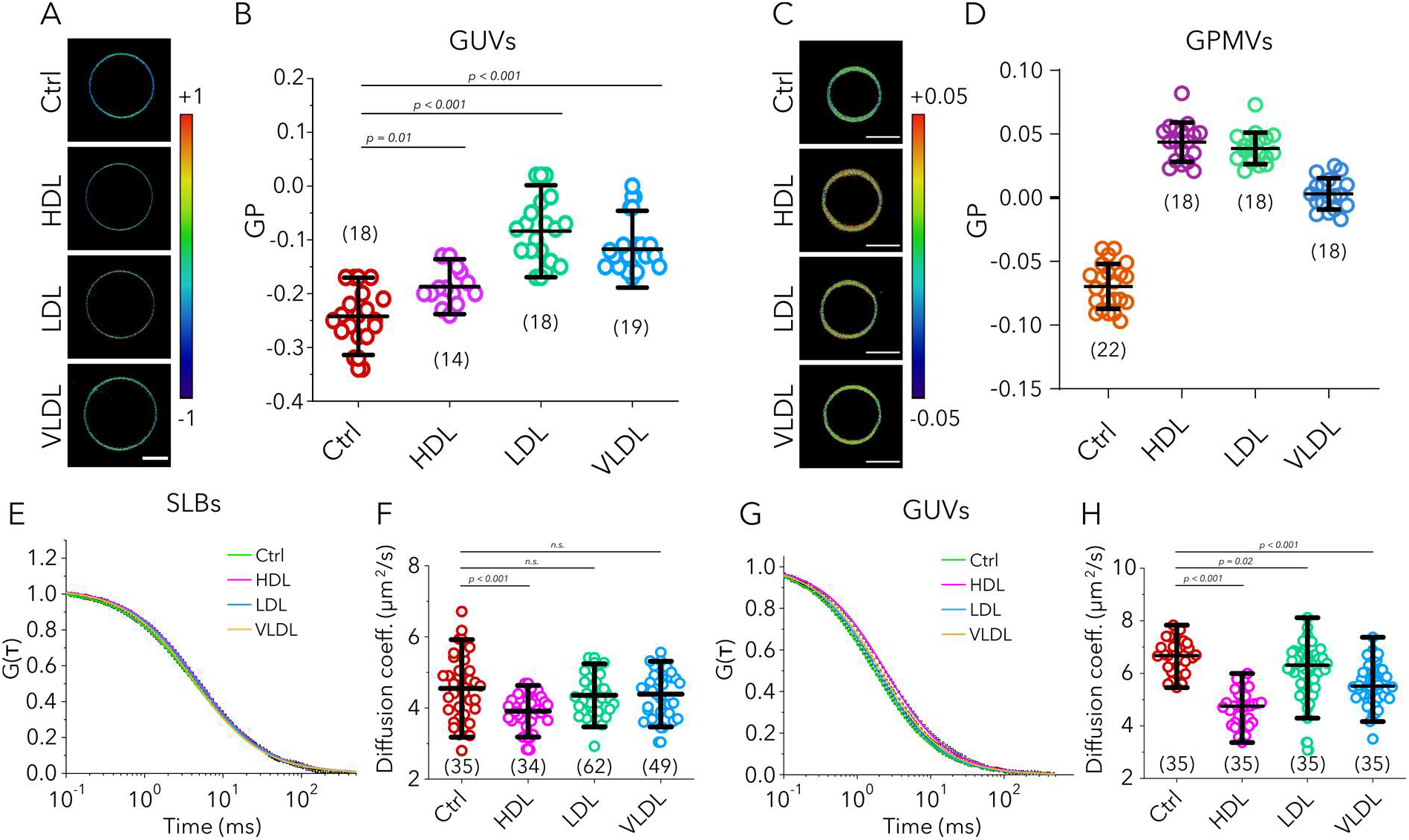
Changes in rigidity upon lipoprotein particle interactions with target membranes. A) GP images and B) GP values of GUVs incubated with HDL, LDL and VLDL particles compared to control GUVs (no incubation). C) GP images and D) GP values of GPMVs incubated with HDL, LDL and VLDL particles compared to control GPMVs. E) Representative FCS curves and F) diffusion coefficients for ASR-PE in SLBs treated with lipoprotein particles compared to control SLBs. G) Representative FCS curves and H) diffusion coefficients for ASR-PE in GUVs treated with lipoprotein particles compared to in untreated GUVs. (Graphs show mean and the standard deviation; number of data points are indicated in graphs in parenthesis)

To further confirm this, we also measured the diffusion of fluorescently-labelled lipid analog (Abberior Star Red-labelled DPPE; ASR-PE) in the membranes of SLBs and GUVs before and after incubation with unlabeled lipoprotein particles with FCS. Diffusion of lipids in rigid membrane is slower compared to more fluid membrane (25). If cholesterol is indeed transferred to the target membrane, it would get more rigid and diffusion would get slower. We observe this trend with all lipoprotein particles (Fig. 3E-H); diffusion coefficient of ASR-PE was 4.5±1.4 μm^2^/s in untreated SLBs while in lipoprotein-treated vesicles it was 3.9±0.7 μm^2^/s (HDL), 4.4±0.8 μm^2^/s (LDL) and 4.4±0.9 μm^2^/s (VLDL) (Fig. 3E, F). In SLBs, the manifestation of compositional changes in diffusion is largely masked by the support effect (25), thus the differences are very small. In contrary, free-standing membranes reflect the compositional changes better (25), thus, we also tested the diffusion of lipid analog in GUVs. In GUVs, diffusion coefficient of ASR-PE was 6.6±1.2 μm^2^/s in untreated GUVs while in lipoprotein-treated vesicles it was 4.7±1.2 μm^2^/s (HDL), 6.1±2.1 μm^2^/s (LDL) and 5.6±1.8 μm^2^/s (VLDL) (Fig. 3G, H). This data together with GP measurements show that lipoprotein particles transfer cholesterol to target membranes and thus increase their rigidity.

## Conclusions

In this work, we showed that all major types of lipoprotein particles (HDL, LDL and VLDL), regardless of their size and their lipid and protein composition integrate with lipid membranes and transfer their shell-derived amphiphilic lipid cargo (exemplified by Bd-Chol) after integration. This adds a new aspect to the picture of cholesterol homeostasis where direct cargo transfer of lipoprotein particles might occur in a receptor-independent manner. This will be the first steps of further work to elucidate the exact contribution of the direct delivery mechanism *in vivo*. Particularly, the efficiency of receptor-independent cholesterol transfer compared to receptor-mediated transfer will shed new light on the relevance of receptor-independent transfer. Despite many unknowns, the ability of the lipoprotein particles to directly deliver their cargo to the target membrane can still potentially be exploited for therapeutic approaches against diseases such as familial hypercholesterolemia where LDL-receptor cannot fulfil its function. Therefore, we believe this mechanism may potentially be important for future therapies against dyslipidemia diseases.

Elaborated work with cells as well as lipoprotein particles from dyslipidemia patients will also be crucial to see how receptor-independent cargo transfer is affected by metabolic state of the donors. Finally, it will be crucial to elucidate whether the target membrane properties influence the lipoprotein interactions, particularly as a function of lipoprotein type.

## Acknowledgements

ES is funded by SciLifeLab and Wellcome Trust Institutional Strategic Support Fund (ISSF). This work has been supported by Austrian Science Fund Project P22838-B13 & P29110-B21 and the Austrian Research Promotion Agency Innovatives Oberösterreich 2020-851455, by the European Fund for Regional Development (EFRE, IWB 2020), the Federal State of Upper Austria, and “Land OÖ Basisfinanzierung”. We acknowledge funding by the Wolfson Foundation, the Medical Research Council (MRC, grant number MC_UU_12010/unit programmes G0902418 and MC_UU_12025), MRC/BBSRC/EPSRC (grant number MR/K01577X/1), the Wellcome Trust (grant ref 104924/14/Z/14). We thank Dr. Andrey Klymchenko for providing NR12S.

## Author Contributions

All authors contributed to experiments, analysis and writing.

