## Supplementary Information for "Lipoprotein particles interact with membranes and transfer their cargo without receptors"

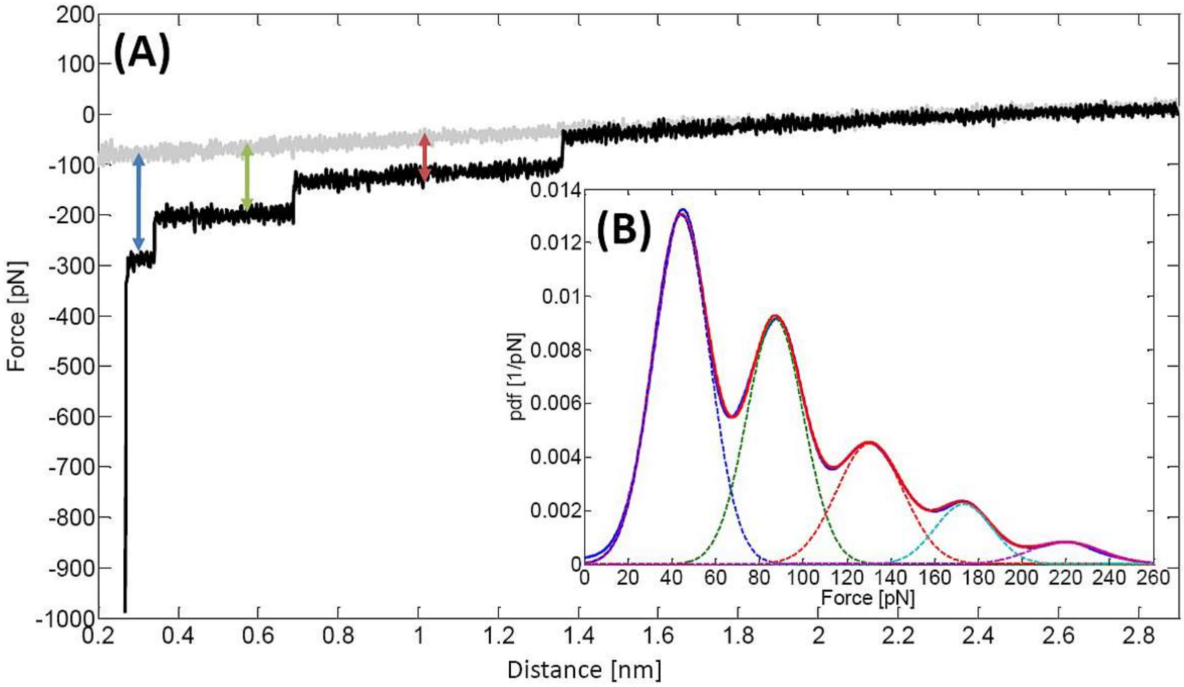


**Figure S1.** Evaluation of the force steps. A) Represents a force-distance cycle, by assuming three force steps. The force for each step was determined between the respective plateau (blue-first step; green-second step; red-second step) and the baseline (grey-approach curve). B) Probability density function of measured forces: The resulting value $\mu_i$ describes the force for the respective step, which was calculated with the provided formula. The plot shows the summarized data of 500 force distance cycles.

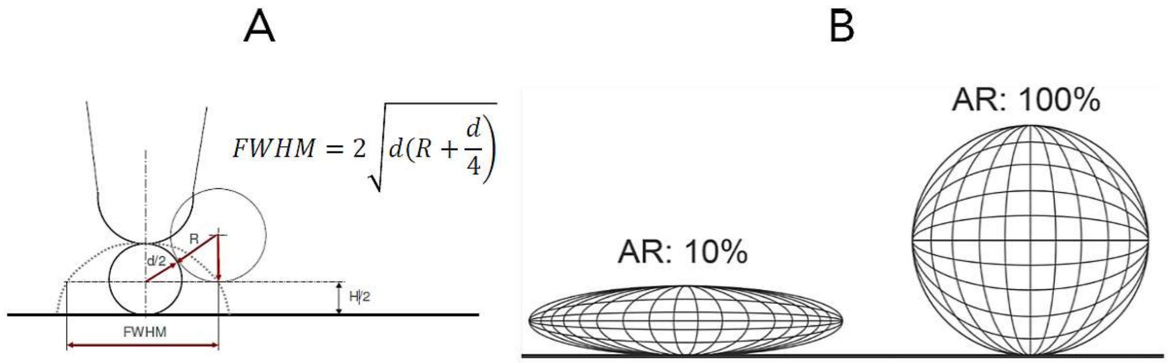


**Figure S2.** (A) Model for the AFM tip convolution: The dotted line represents the height information of a spherical object, which is detected by an AFM tip if their diameters are nearly identical (diameter AFM tip apex (R) ~ diameter spherical particle (d)). By measuring the Full Width at Half Maximum (FWHM) from the AFM images and solving the equation (see Figure S1 (A)) for the parameter d, the deconvoluted particle width is obtained. (B) Two representations of particles with an Aspect Ratio (AR=Height/Width *100%) of 10% and 100%.

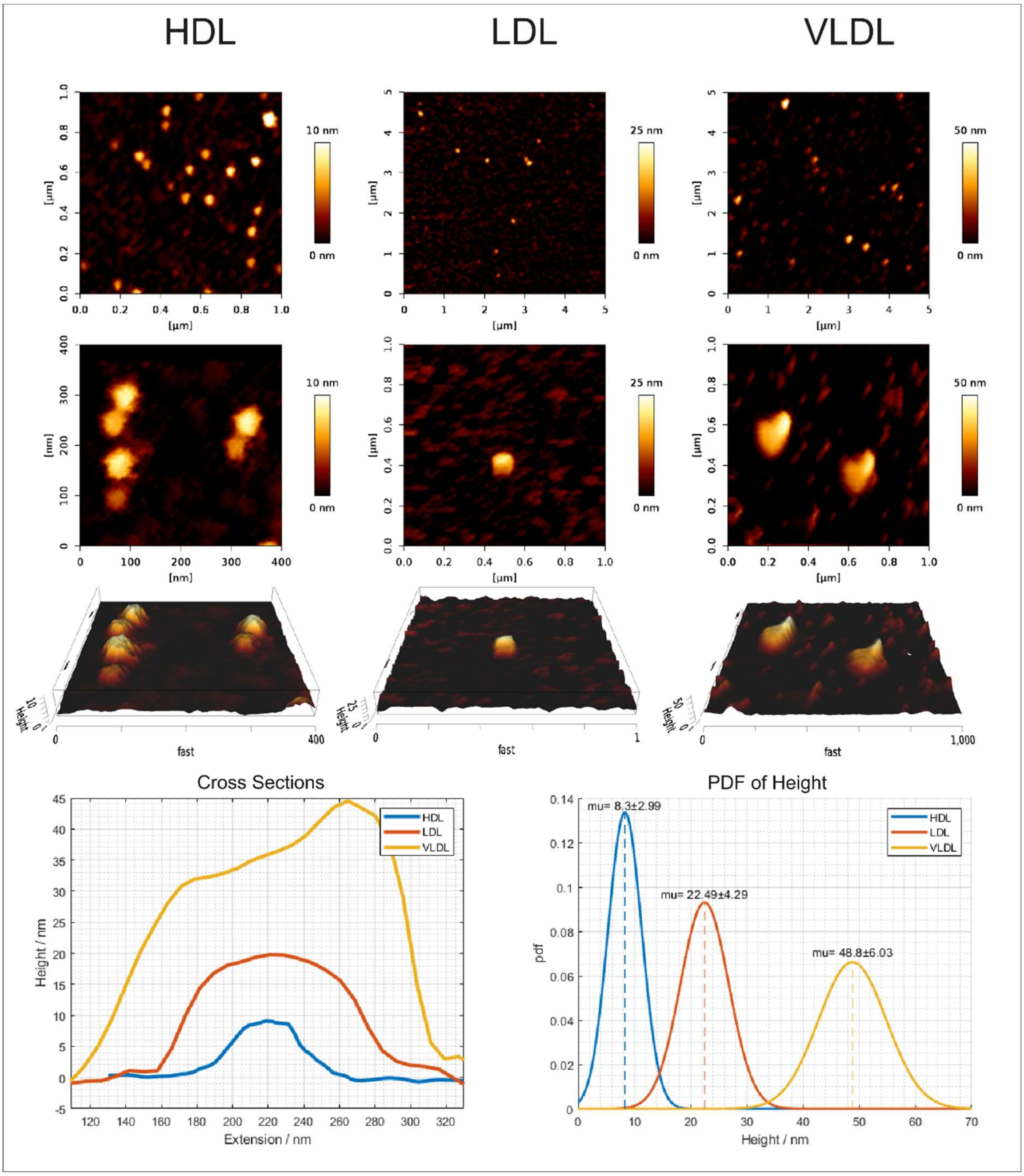


**Figure S3.** AFM height image overview (first row), increased magnification (second row) and 3D display of magnified HDL, LDL and VLDL particles (third row) on adsorbed on glass cover slip. Cross section figure of magnified HDL, LDL, and VLDL particles. Probability Density Function (PDF) of analyzed HDL, LDL and VLDL particle heights. The population average of the PDF functions of the respective particles are: HDL=8.30±2.99 nm, LDL=22.49±4.29 nm and VLDL=48.80±6.03 nm (μ±σ) n=10.

Membrane nanotubes (or tethers) are nanocylinders of lipid bilayers, arising from manipulation experiments on synthetic vesicles, to formation of dynamic tubular networks in cells. However, here multiple tether formation was observed on a supported lipid bilayer, consisting only of DOPC. In particular, the average force required to form a single tether was found to be ~50pN, at different cantilever retraction speed between 0.5 – 5 µm/s (Fig. S4C). Typically, forces were calculated by fitting a Gaussian profile into the force pdf (see Fig. S4A, B). By plotting the force versus the length of the tethers, well defined equidistant force steps were extracted (Fig. S4C). Up to four unbinding events were observed, indicating multiple tether formation (see Fig. S4D).

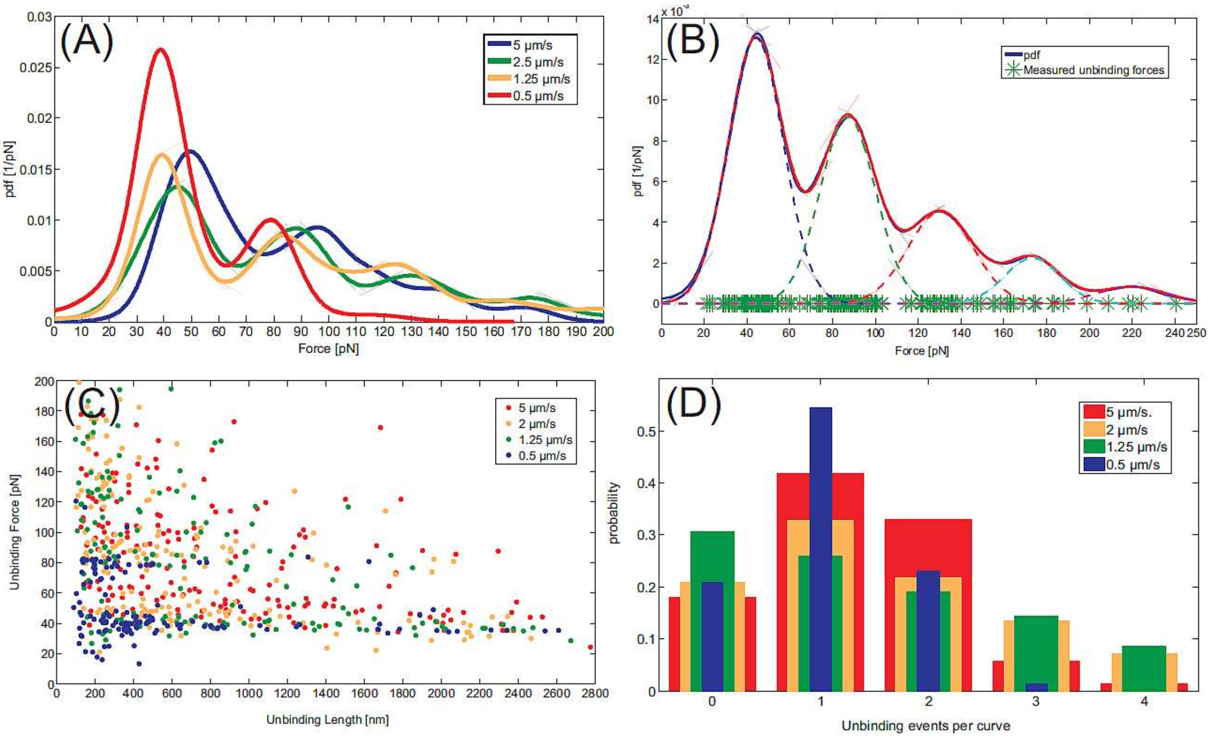


**Figure S4.** Dependence of force of individual steps on the pulling velocity. The retraction velocity ranges from 0.5-5 µm/s. Colors are consistent for all images: red 0.5 µm/s, yellow 1.25 µm/s, green 2.5 µm/s, clue 5 µm/s. A) Force pdf for varying retraction speeds B) The exact forces were extracted through a Gaussian Fit C) Force versus length: Independent on the adjusted pulling velocity well defined force steps were measured D) Probability of unbinding events: Up to four unbinding events were detected.

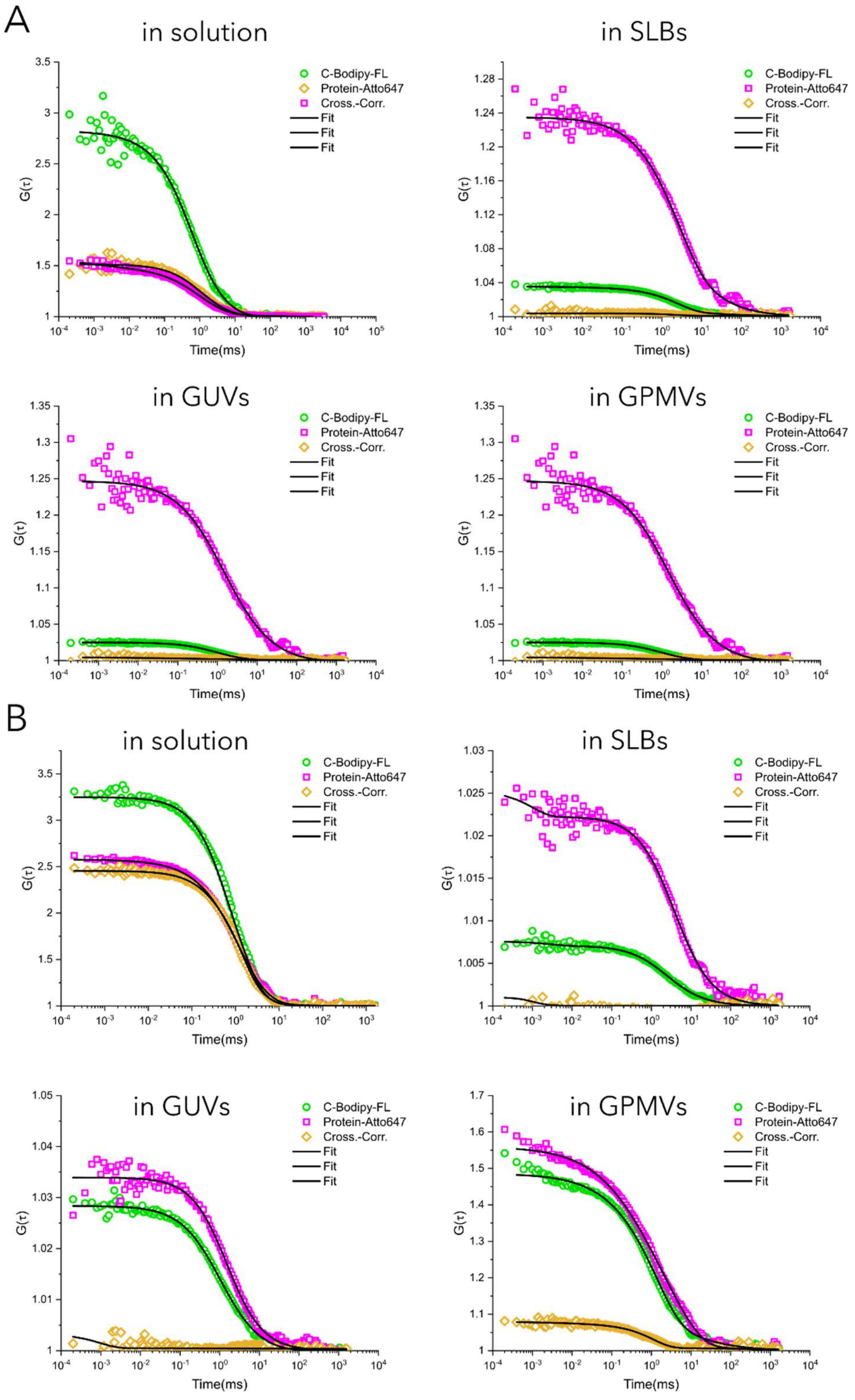


**Figure S5.** Fluorescence cross-correlation spectroscopy of C-BodipyFL and proteins measured in solution (intact lipoproteins) as well as in target membranes (GUVs, GPMVs and SLBs – as labeled) for A) HDL and B) LDL. In solution, high cross-correlation of C-BodipyFL and protein signals is detected which suggests co-diffusion. In target membrane, cross-correlation curve amplitude is close to zero which suggests that C-BodipyFL and protein molecules diffuse in the target membrane independently.

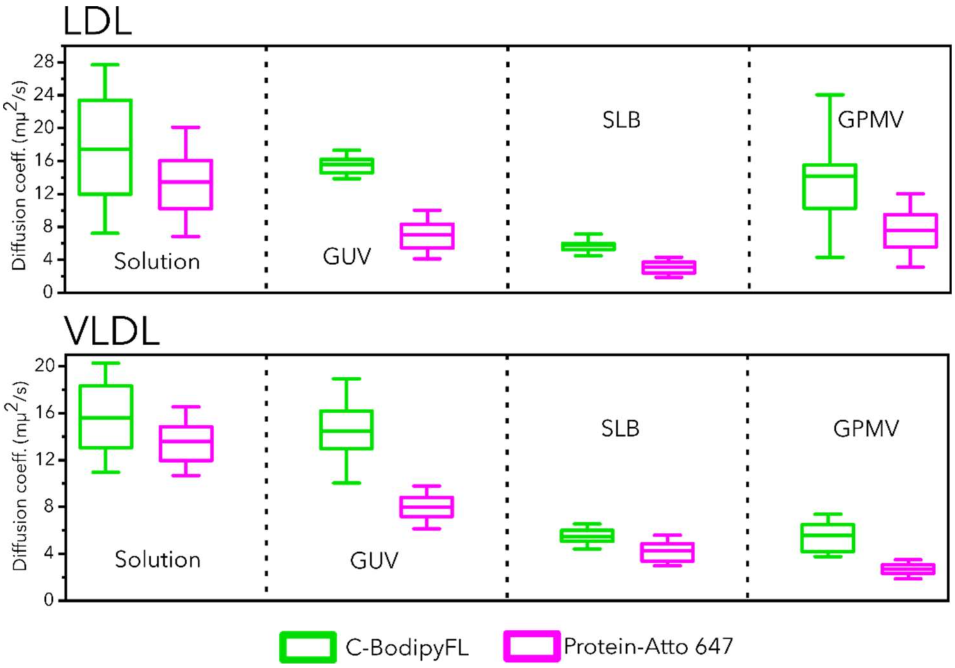


**Figure S6**: Diffusion coefficients of C-BodipyFL and proteins measured in solution (intact lipoproteins) as well as in target membranes (GUVs, GPMVs and SLBs – as labeled) for A) HDL and B) LDL. In solution, the diffusion coefficients of C-BodipyFL and proteins are similar, because they diffuse together as intact lipoprotein. In all target membranes, C-BodipyFL diffuses faster as protein molecules.
